# Transcriptional Repression by RelA and Yin Yang1 is essential for survival of Colorectal Cancer Cells

**DOI:** 10.1101/2025.05.12.653423

**Authors:** Ramadevi Mutra, Sivakumar Vallabhapurapu

**Affiliations:** Indian Institute of Science Education and Research Tirupati, Andhra Pradesh, India

## Abstract

NF-κB is primarily known for its transcriptional activation function in the context of immune responses, inflammation, cell survival, proliferation and Cancer. However, whether NF-κB functions as a transcriptional repressor in the context of tumor growth has not been well addressed. While, classical NF-kB activated by IKK-b has been shown to play a tumor promoting role in colorectal cancer, couple of studies suggested that overexpression of RelA in colorectal cancer cells leads to apoptosis. These findings were contradictor and puzzling with regards to the role of RelA in colorectal cancer. Here, we report that RelA represses proapoptotic gene puma and contributes to colorectal cancer cell survival. Interestingly, Yin-Yang1, a RelA target gene product is also essential to repress puma and contributes to colorectal cancer cell survival. Moreover, RelA and YY1 form a complex in colorectal cancer cells. Depletion of either RelA or YY1 results in upregulation of pro-apoptotic gene Puma. Importantly, we show that binding of Yin Yang1 to RelA impairs transcriptional activation by RelA suggesting that YY1 is an inhibitor of RelA function. Collectively, we present evidence for RelA-YY1 complex formation in colorectal cancer cells, both RelA and YY1 function as transcriptional repressors of puma and contribute to survival of colorectal cancer cells.

## Introduction

Among different cancers that affect human health, colorectal cancer (CRC) ranks within the top three in causing cancer-related deaths (1,2). Several new cases of colorectal cancers are diagnosed each year world wide. (3). While several oncogenic factors including the JAK-STAT, Ras, Akt and the mTOR (4-6) pathways are known to be involved in colorectal cancer progression, it is of importance to note that chronic inflammation activating the NF-κB pathway has been shown as a major risk factor in the progression of CRC (7, 8). While the inflammatory microenvironment created by IKKβ that activates the RelA subunit within CRC tumor tissue has been shown to be essential for CRC growth, the precise mechanism by which RelA contributes to CRC growth is not fully understood. Interestingly, one of the RelA target genes Yin-Yang1 (YY1) is also highly expressed in CRC and contributes to CRC progression (9). However, how RelA and YY1 coordinate and contribute to CRC progression has not been addressed. Further, while patients diagnosed with early stage CRC have better prognosis, those diagnosed with advanced stage CRC exhibit poor prognosis as it is difficult to treat advanced CRC (10). Hence, identification of novel gene regulatory complexes that play a role in survival of advanced stage CRC cells will be helpful in designing novel and targeted therapeutic approaches for CRC treatment.

In mammals, different NF-κB family of transcription factors including RelA, c-Rel, RelB, NF-κB2 and NF-κB1 are regulated differently by distinct upstream kinases and signal transduction pathways (11-13). Of these, NF-κB2 and NF-κB1 encode for the full length p100 and p105 respectively, which eventually get processed to produce p52 and p50 (11-13). These factors are known to form different homo and heterodimers that are activated by classical or the alternative NF-κB activation pathways. While the classical NF-κB pathway is regulated by the IKK complex comprising IKKβ, IKKα and IKKγ subunits and contribute to the activation of p50-RelA complex, the alternative pathway is regulated by the kinase NIK and IKKα to activate p52-RelB complex (11-14). Once, in the nucleus, the classical and alternative NF-κB complexes are primarily known to function as transcriptional activators to regulate distinct sets of target genes that play diverse roles in immune responses, inflammation, cell survival and proliferation as well as tumor progression (15). While some work from the Perkins group has shed early light into potential transcriptional repressive role by NF-κB (16), little is known whether NF-κB functions as transcriptional repressor to promote tumor growth. Moreover, although NF-κB is known to be frequently activated in most cancers, the mechanism by which classical and the alternative NF-κB complexes contribute to cancer progression is expected to be different in different cancers (17-19). Interestingly, in CRC, although classical NF-κB was found to be constitutively activated (20), exogenous overexpression of RelA and phospho-mimic (at S536) RelA was found to enhance apoptosis of CRC cells (21,22). This raises a question as to whether RelA is essential for CRC cell survival and whether RelA acts as a pro-tumorigenic factor or tumor suppressor in CRC. Moreover, while NF-κB is often activated in several cancers, it also plays important role for normal homeostasis and hence the use of existing NF-κB inhibitors targeting IKK or NIK would cause severe toxic side effects (11). Hence, identification of novel NF-κB complexes that are essential for tumor growth would enable us to develop more targeted therapeutic approaches than blocking IKK / NIK that would block the entire pathway.

Yin-Yang1 (YY1) is a dual function transcription factor that functions both as a transcriptional activator and repressor and has also been found to be hyper expressed in several tumors including CRC (9). While YY1 was found to act both as a tumor promoter and tumor suppressor, in CRC, YY1 has been linked to tumor progression (9, 23) by functioning as a transcriptional activator inducing the expression of diverse protumorigenic genes (9). However, whether YY1 acts as a transcriptional repressor in CRC progression and importantly, whether YY1 coordinates with NF-κB subunit RelA in CRC progression has not been addressed.

We have previously reported that RelA and YY1 interact to form a transcriptional repressive RelA-YY1 complex to repress the proapoptotic gene Bim and promote multiple myeloma cell survival (24). Here we report that RelA and YY1 interact to form a complex in CRC cells and both RelA and YY1 are essential for CRC cell survival by repressing a pro-apoptotic gene PUMA. Importantly, we report that YY1 binds to RelA and inhibits the transcriptional activation by RelA.

## Results

### RelA and YY1 are essential for CRC cell survival

It was previously shown that RelA overexpression in CRC cells enhances their apoptosis (21,22) raising a question about the actual role of RelA in CRC cell survival. Here we show that depletion of RelA by lentiviral mediated expression of RelA specific Sh-RNA causes apoptosis of CRC cells (Fig. 1A – 1C). Like wise, depletion of YY1 by lentivirally expressed YY1 specific Sh-RNA causes CRC cell apoptosis (Fig. 2A – 2C) which is in line with previous reports (23).

**Figure 1.**
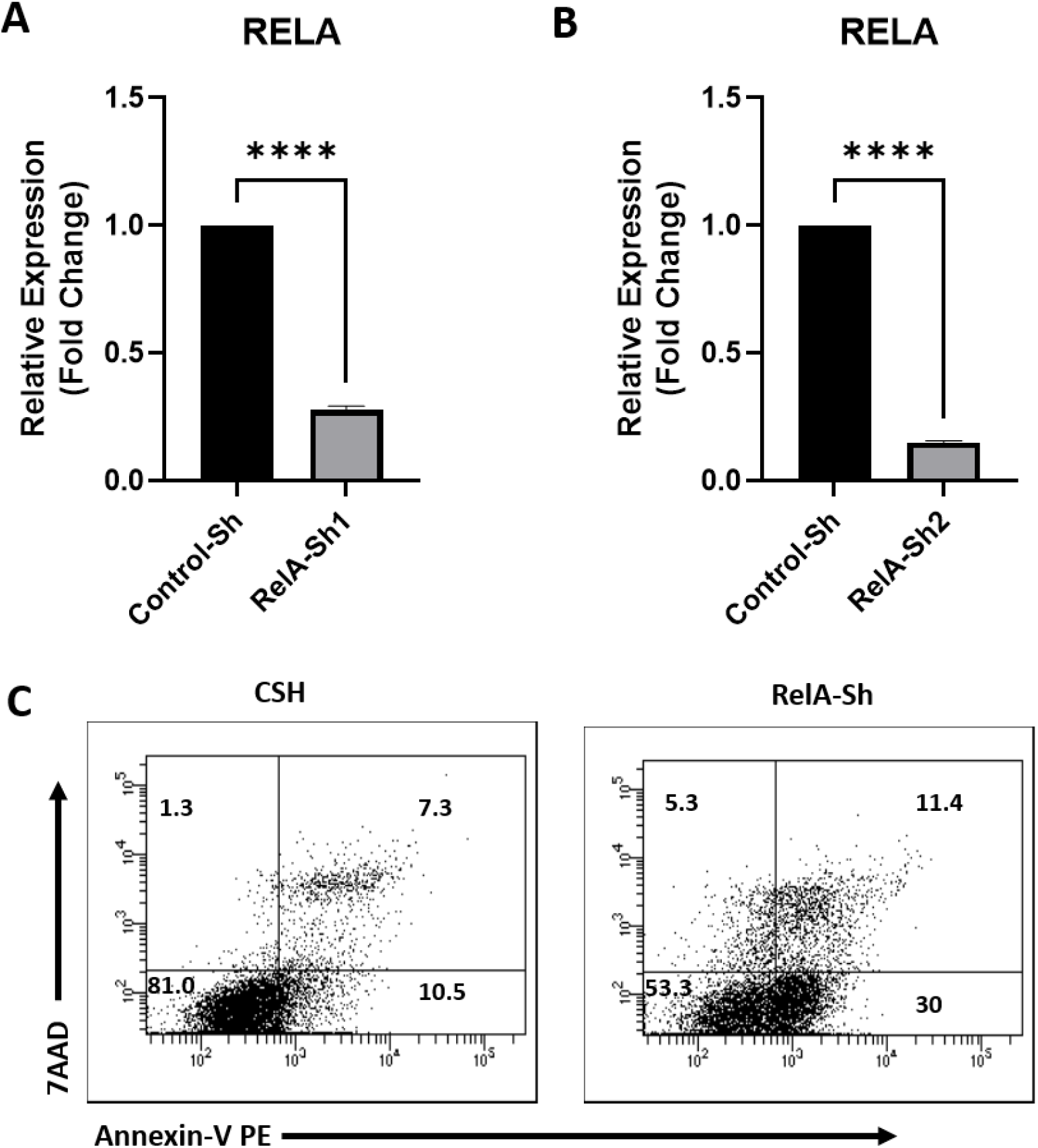
RelA is essential for CRC cell survival: A and B) HCT-116 cells were tranduced with lentiviral vector expressing RelA-specific Sh-RNA. Cells were lysed and total RNA was collected 48 hours after transduction and RelA-knockdown was analyzed by real-time PCR analysis. C) HCT-116 cells were transduced with lentiviral vector expressing a control non-targetting sh-RNA or RelA-specific Sh-RNA. 4 days after transduction cells were analyzed for survival by flow cytometry upon Annexin-V and 7AAD staining. Numbers in the plot indicate relative percentages of cells.

**Figure 2.**
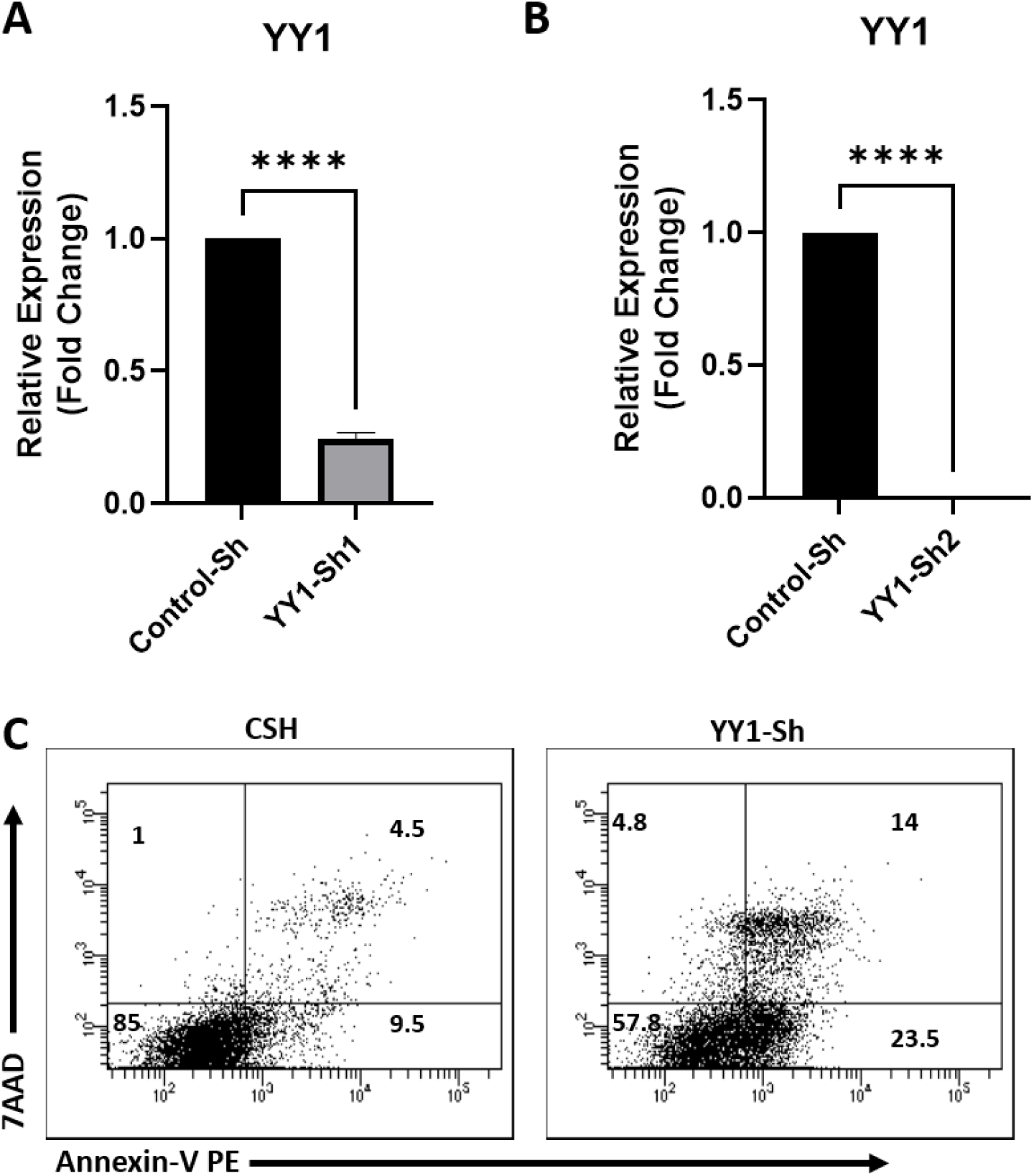
YY1 is essential for CRC cell survival. A and B) HCT-116 cells were tranduced with lentiviral vector expressing YY1-specific Sh-RNA. Cells were lysed and total RNA was collected 48 hours after transduction and YY1-knockdown was analyzed by real-time PCR analysis. C) HCT-116 cells were transduced with lentiviral vector expressing a control non-targetting sh-RNA or YY1-specific Sh-RNA. 4 days after transduction cells were analyzed for survival by flow cytometry upon Annexin-V and 7AAD staining. Numbers in the plot indicate relative percentages of cells.

### Both RelA and YY1 are essential to repress PUMA

We show that mechanistically, RelA and YY1 both contribute to CRC cell survival by repressing the expression of proapoptotic gene puma. As shown in Fig. 3A and 3B, depletion of RelA by lentiviral expression of RelA specific Sh-RNAs in HCT116 cells resulted in the increased expression of puma both at mRNA and protein levels. Likewise, lentiviral expression of YY1-specific ShRNAs resulted in elevated levels of puma mRNA and protein (Fig. 4A and 4B). However, we did not observe significant changes in the expression of other pro-apoptotic genes upon depletion of RelA or YY1 (data not shown). In line with these observations, by employing luciferase reporter assay using pGL2-puma reporter vector, we found that both RelA and YY1 could significantly repress puma promoter activity (Fig. 5). These results suggest that transcriptional repression of puma by both RelA and YY1 is essential for CRC cell survival.

**Figure 3.**
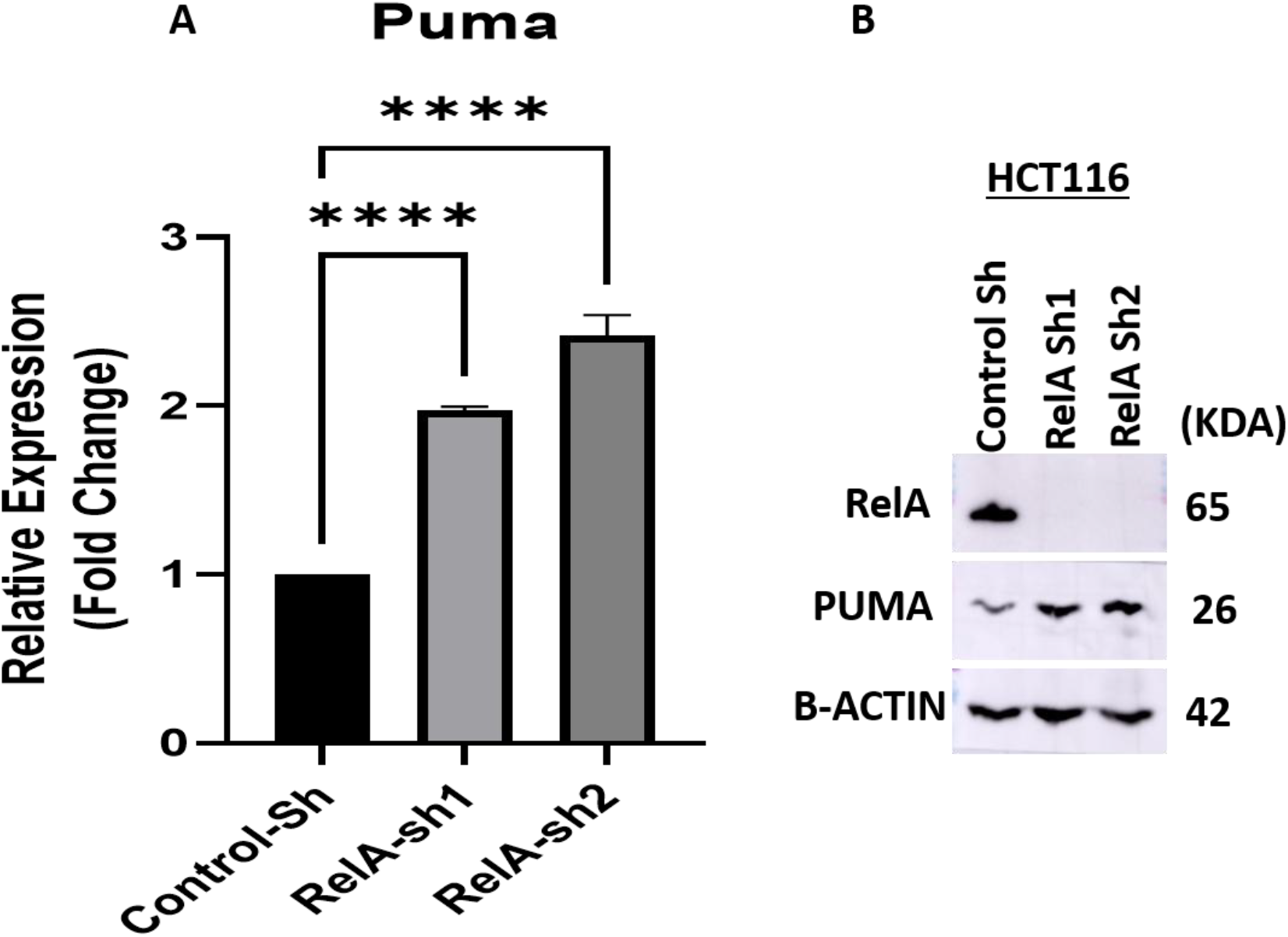
RelA represses puma in colorectal cancer cells. A) Total RNA from control and RelA-depleted cells was isolated, reverse transcribed and analyzed for the expression of puma by real time pcr. B) Whole cell lysates from control and RelA-depleted cells were isolated and immnoblotted and puma expression was analyzed as indicated.

**Figure 4.**
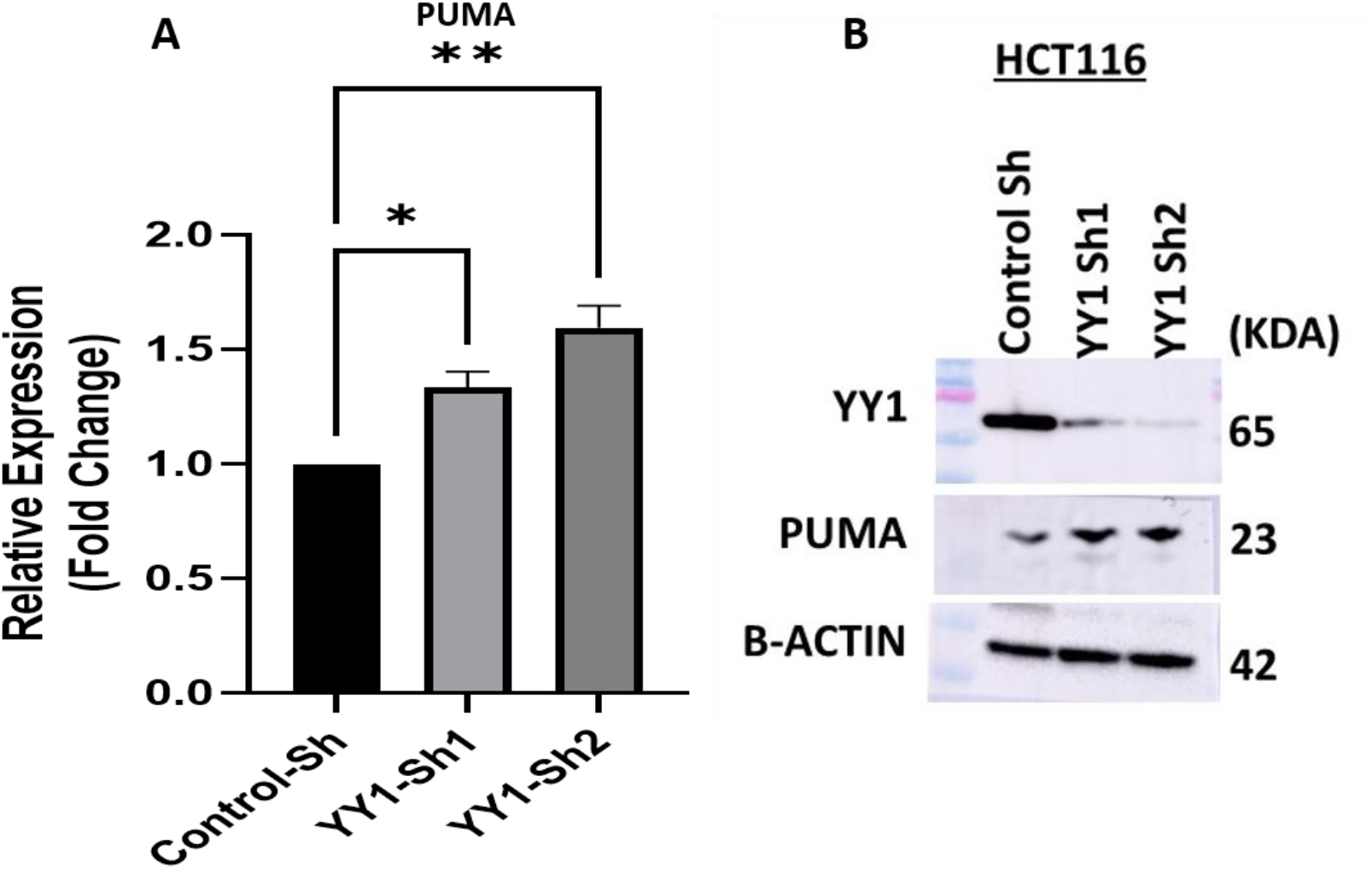
YY1 represses puma in colorectal cancer cells. A) Total RNA from control and YY1-depleted cells was isolated, reverse transcribed and analyzed for the expression of puma by real time pcr. B) Whole cell lysates from control and YY1-depleted cells were isolated and immnoblotted and puma expression was analyzed as indicated.

**Figure 5.**
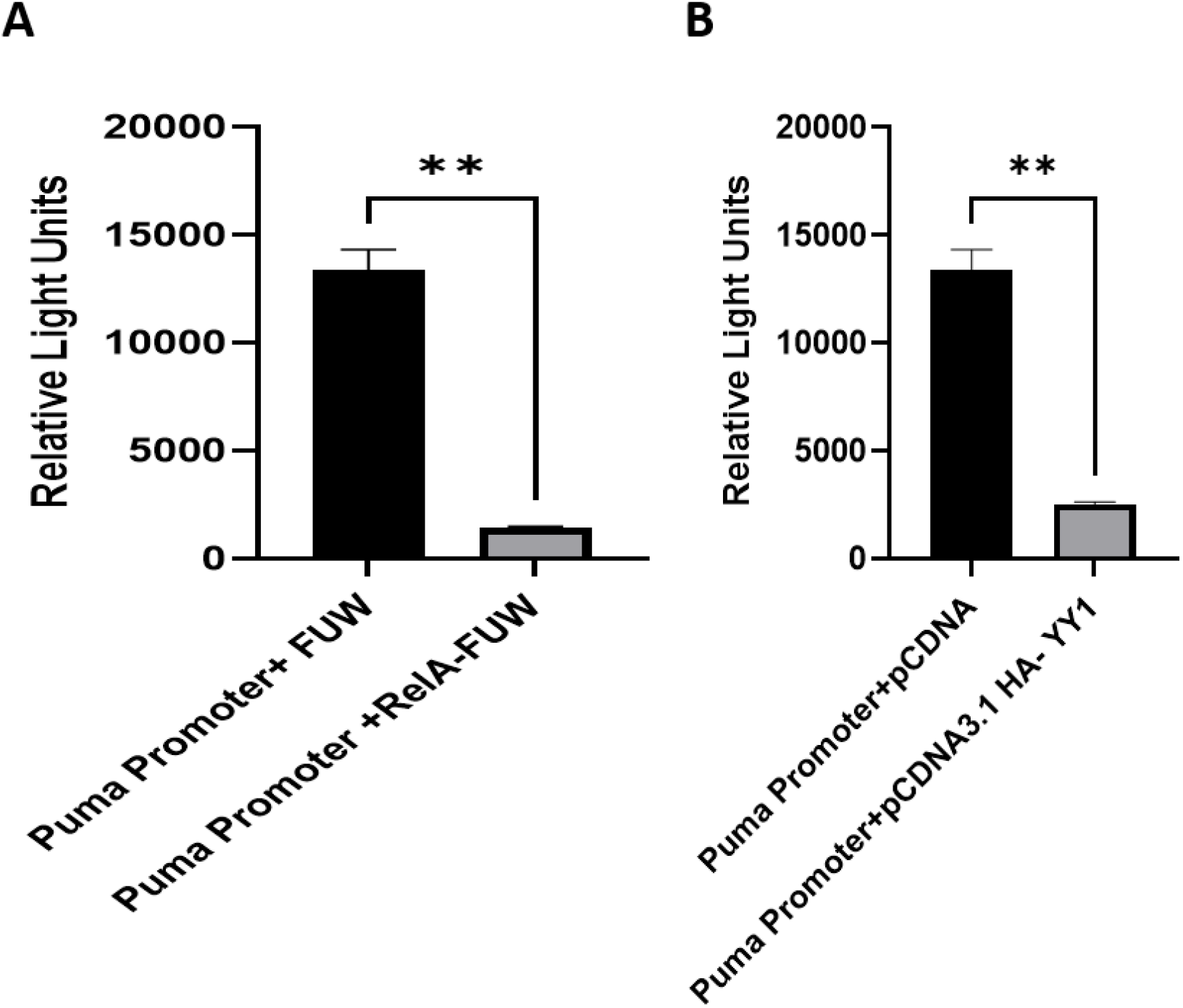
Both RelA and YY1 repress puma promoter. A) HEK-293T cells were transfected with pgl2-puma luciferase reporter vector alone or in combination with RelA or YY1. Relative luciferase activity was shown as light units observed.

### RelA and YY1 interact to form a complex in CRC

Previously we have shown that RelA and YY1 interact with each other to form RelA-YY1 complex that repress proapoptotic gene bim in multiple myeloma cells (24). However, it is not clear whether RelA and YY1 form a complex in other cancer models including CRC. Since both RelA and YY1 were found to be essential to repress puma in HCT116 cells, we tested whether RelA and YY1 physically interact to form RelA-YY1 complex in CRC cells. To this end, we employed immunoprecipitation experiments in HCT116 cells and found that YY1 can be immunoprecipitated by RelA and vice versa as shown in Fig. 6 indicating constitutive RelA-YY1 complex formation in CRC cells.

**Figure 6.**
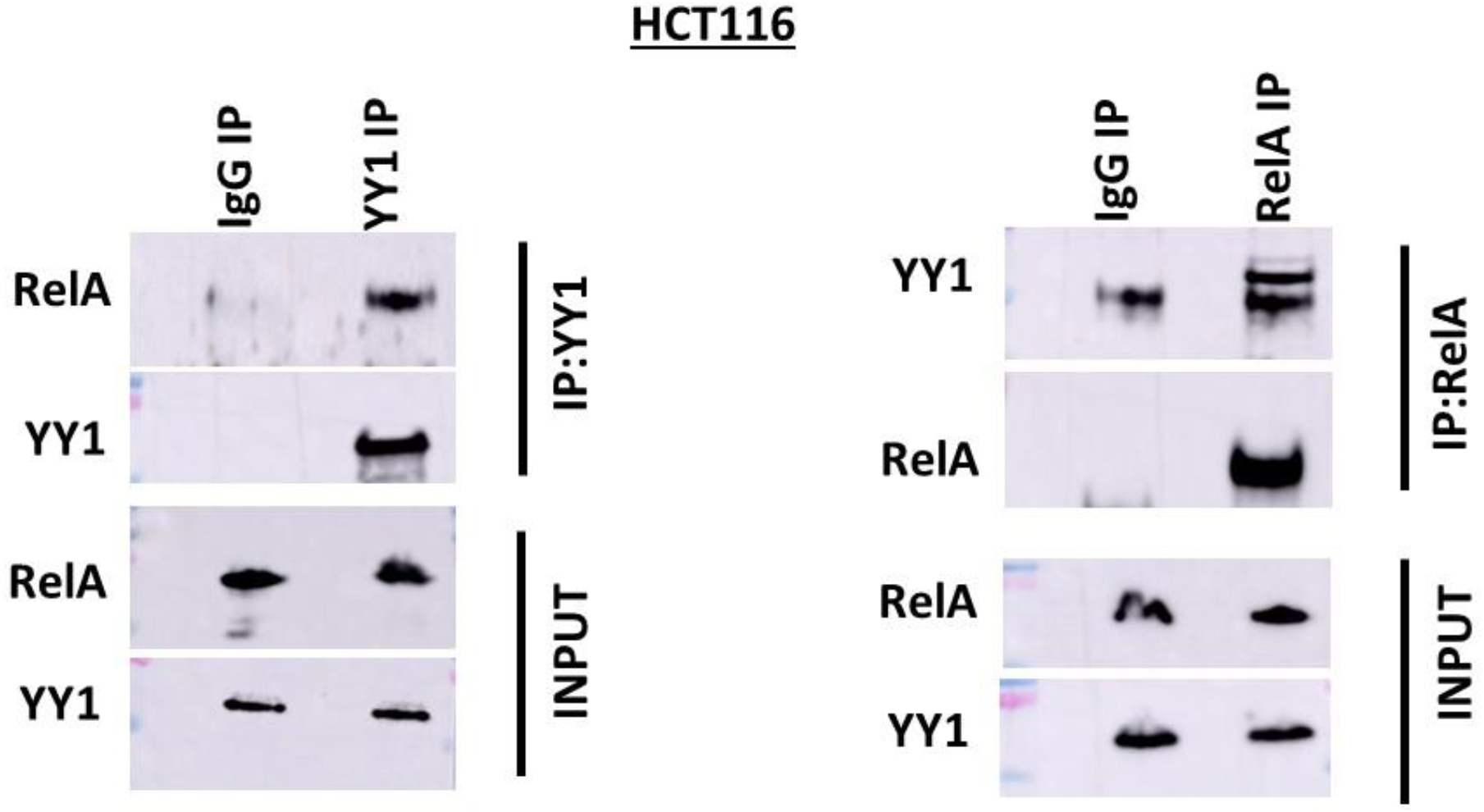
RelA and YY1 interact to form a complex in colorectal cancer cells. HCT116 cells were lysed and both RelA and YY1 were separately immunoprecipitated and the immunoprecipitated complexes were immunoblotted with indicated proteins. Note the interaction of RelA and YY1.

### YY1 is an inhibitor of transcriptional activation by RelA

While we showed that RelA-YY1 complex functions as a transcriptional repression on pro-apoptotic genes (24), it is not clear whether YY1 interaction with RelA would interfere with RelA-mediated transcription of its target genes. To this end, we performed luciferase reporter assays by co-transfecting a known RelA reporter plasmid containing 3X-κB RelA binding sites along with RelA and confirmed that RelA significantly induces luciferase expression as expected. However, co-transfecting YY1 along with RelA and 3X-κB reporter plasmid has significantly reduced luciferase expression clearly indicating that YY1 impairs transcriptional activation function of RelA (Fig. 7A – 7C). To further test whether constitutive binding of YY1 to RelA will result in loss of transcriptional activation of by RelA, we made a RelA-hinge-YY1 fusion construct, in which RelA and YY1 were fused to each other with a immunoglobulin hinge region between them (Fig. 7A and 7B). Interestingly, co-transfection of RelA-hinge-YY1 fusion construct (Fig. 7B and 7C) with 3X-κB reporter plasmid resulted in almost no expression of luciferase indicating complete loss of RelA-dependent transcription upon constitutive binding to YY1. Thus, these results indicate that YY1 is a direct inhibitor transcriptional activation by RelA.

**Figure 7.**
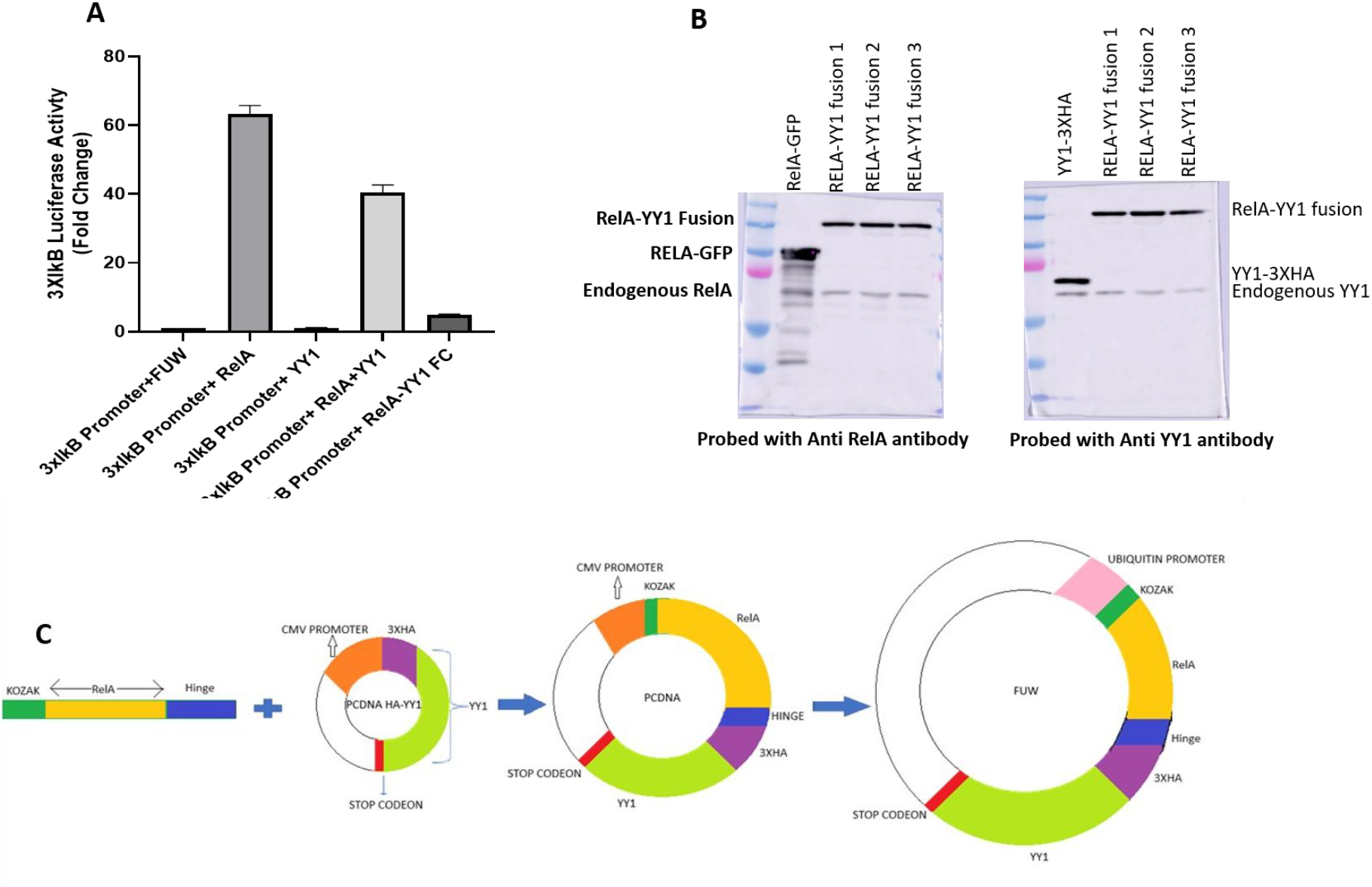
YY1 inhibits RelA-dependent transcription. (A) HEK-293T cells were transfected with a 3X-kb reporter plasmid (NF-kB reporter) either alone or in combination with RelA, RelA + YY1 or with RelA-YY1 fusion construct (B and C). Relative luciferase activity was shown as light units in the form of a bar diagram. Also shown in the figure is a schematic diagram for RelA-hinge-YY1 fusion construct design (C). HEK-293T cells were transfected with the RelA-hinge-YY1 construct (3 different clones) and whole cell lysates prepared were analyzed by immunoblotting for indicated proteins to confirm the expression of RelA-hinge-YY1 fusion protein (B).

## Discussion

Colorectal Cancer (CRC) is a major health problem across the globe and several oncogenic factors have been identified to be essential for CRC progression. While diverse classes of drugs are known to be employed in CRC therapy, majority of them cause toxic side effects and complete cure for CRC is still not feasible. Hence, new drugs targeting novel oncogenic factors in CRC are needed for better therapy and less toxic side effects. Among the oncogenic factors known to be involved in CRC progression, NF-κB pathway activated by IKKβ has been shown to play key role (7). IKKβ has been shown to largely activate the classical NF-κB (RelA containing complexes) pathway. Moreover, IKKβ is a major kinase that phosphorylate RelA at serine 536, which enhances the transcriptional activation by RelA (11, 25). This suggests that IKKβ driven activation of RelA in CRC might be a key oncogenic factor. However, surprisingly, it was reported that overexpression of RelA or phospho-mimic (S-536) RelA was shown to be toxic to CRC and reduce CRC growth (21.22). Moreover, treatment with TNF-a was shown to cause CRC cell death via NF-kB (RelA) mediated induction of the pro-apoptotic gene puma (26). These results were mutually contradictory and makes it puzzling with regards to whether RelA is pro-tumorigenic or tumor suppressor in CRC.. To address this directly, we depleted RelA in CRC cells by employing lentiviral mediated expression of Sh-RNA specific to RelA and found that RelA depletion causes significant apoptosis of CRC cells. Mechanistially, we showed that RelA represses the proapoptotic gene puma in CRC cells and thereby contribute to CRC cell survival. Moreover, we showed evidence that RelA represses puma gene prmoter. This indicate that while RelA is certainly required for CRC survival, more detailed studies are needed to address the role of S-536 phosphorylated RelA in CRC.

Yin-Yang1 is a dual function transcription factor with roles both as transcriptional activator and repressor (9). Likewise, YY1 has also been shown to function both as a tumor promotor and tumor suppressor (9). However, in CRC, YY1 has been majorly shown to act as tumor promoter by functioning as a transcriptional activator (9) and it was not clear whether transcriptional repression by YY1 plays a role in CRC progression. Here, we show that in CRC cells YY1 is essential for repression of the proapoptotic gene puma and thereby contribute to the survival of CRC cells as shown by the increased apoptosis of CRC cells upon YY1 depletion. Importantly, we show here that YY1 readily forms a complex with RelA and the function of RelA-YY1 complex in CRC cells needs to be studied further. We have previously shown that RelA-YY1 complex represses another proapoptotic gene bim in multiple myeloma cells (24). Our results presented here document that both RelA and YY1 are essential for repression of puma and that RelA and YY1 interact with each other to form a complex. Hence, it is possible that RelA-YY1 as a complex is essential for puma repression and CRC cell survival. Importantly, while we found that RelA-YY1 complex is detected in HCT-116 cells, we did not observe this complex in HT-29 cells, suggesting that RelA-YY1 complex might be present in more aggressive CRC cells perhaps with cells harboring KRAS mutations. Hence, RelA-YY1 targetting strategies might be helpful to develop novel therapeutic methods for aggressive CRC treatment.

While RelA-YY1 complex appear to function as transcriptional repressor of Bim, it was not clear what are the physiological roles of this complex. Our data presented here indicate that YY1 binding to RelA interferes with its transcriptional activation function (Figure 7). Importantly, our results obtained with RelA-Hinge-YY1 fusion construct indicate that constutitive binding of YY1 to RelA severely impairs transcriptional activation by RelA. This is interesting and YY1 mediated inhibition of transcription of RelA needs to be investigated in the immune system and cancer further. Importantly, YY1 has been known to be a target gene of RelA (27). Also, inflammatory signals such as TNF are known to induce YY1 in a RelA dependent manner (27). Hence, whether YY1 mediated inhibition of transcriptional activation by RelA plays a role to downmodulate inflammation during physiological responses would be an interesting aspect to study.

## Material and Methods

### Cell culture

HEK-293T and HCT116 cells were procured from ATCC and were cultured according to ATCC culture conditions. Both Cell lines were cultured in DMEM medium supplemented with 10% FBS and 1% Penicillin-Streptomycin in an incubator maintained at 5% CO2 and 37C.

### Plasmids and transfection for lentiviral production

pLKO.1 puro Lentiviral vectors targeting RelA and YY1 were purchased from Sigma Aldrich. Non-targeting pLKO.1 puro plasmid (CSH), pMD2.G (VSV G) envelope plasmid and the Gag, Pol expressing psPAX2 packaging plasmid were procured from Addgene. Lentiviruses expressing control ShRNA (CSH), RelA-ShRNA (RelA-Sh) and YY1-ShRNA (YY1-Sh) were prepared by co-transfecting pLKO.1 plasmid (for respective Sh-RNAs) along with pMD2.G and psPAX2 in HEK-293T cells. 72 hours post transfection lentiviruses were collected from the supernatants.

### Lentiviral transduction and gene silencing

HCT116 cell lines were seeded in 6 well plates. Once the cells achieved 70% of confluency they were transduced with lentiviruses expressing Control ShRNA (addgene#109012), RelA-specific (TRCN0000014684 and TRCN0000014683) and YY1-specific ShRNAs (TRCN0000019898 and TRCN0000019894) separately. Sixty hours post transduction cells were lysed to collect total RNA and whole cell lysates for further analysis.

### Cell Viability Assay

4 days after lentiviral transduction, all the cells in the plate were collected and viability of cells was analysed by flow cytometry using FACS-Celesta (BD Biosciences), upon staining with Annexin-V and 7AAD (BD Biosciences) according to the manufacturer’s recommendations.

### Immunoblot and qPCR Methods

For whole Cell lysate preparation, cells were lysed in 1x Triton X-100 lysis buffer supplemented with Protease inhibitor (Roche Cat# 11873580001) and phosphatase inhibitor (Roche Cat# 04906845001) cocktail. For immunoprecipitation cells were lysed in IP lysis buffer (Thermo Cat# 87787) supplemented with protease and phosphatase inhibitor cocktail. Immunoprecipitations were performed by incubating whole cell lysates were first precleared by incubation with Protein G dynabeads (Thermo Cat#10004D) for 1 h at 4 °C. Precleared lysates were immunoprecipitated by incubating with specific antibodies at 4 °C overnight and further incubated upon adding Protein G dynabeads for 1 h at 4 °C. The beads were collected using magnetic separator and were washed extensively with IP lysis buffer mentioned above. Immunoprecipitated proteins were then eluted by incubating with 2X SDS sample loading dye at 95°C for 5 min. Whole cell lysates, and immunoprecipitates were separated by SDS–PAGE and analysed by immunoblotting with specific antibodies. Antibodies used were RelA (sc-8008), YY1(Cell Signaling Technology - 46395S), beta tubulin (Cell Signaling Technology-86298S), beta actin (Cloud clone - MAB340Mi21), puma (Cell Signaling Technology-4976s), anti-HA (sc-7392), normal Rabbit IgG (Cell Signaling Technology-2729S), normal Mouse IgG (sc-2025). For gene expression analysis by Quantitative real-time PCR, cells were lysed and total RNA was isolated from using RNeasy Plus kit (Qiagen). We the proceeded to make first strand cDNA by reverse transcription using iScript cDNA synthesis kit (Bio-Rad). Real-time PCR was performed on Bio-Rad (CFX Connect Real Time System). Various primers used were: Bim Forward: 5′-TCAACACAAACCCCAAGTCC -3′, Reverse: 5′-TAACCATTCGTGGGTGGTCT -3′; BMF Forward: 5′-GAGGTACAGATTGCCCGAAA -3′, Reverse: 5′-CCCCGTTCCTGTTCTCTTCT -3′; Bad Forward: 5′-CGAGTGAGCAGGAAGACTCC -3′, Reverse: 5′-ATGATGGCTGCTGCTGGT -3′; BOK Forward: 5′-GGCGATGAGCTGGAGATGA - 3′, Reverse: 5′-ACCGCATACAGGGACACCA -3′; Bid Forward: 5′-GTGAGGTCAACAACGGTTCC -3′, Reverse: 5′-TATTCTTCCCAAGCGGGAGT -3′; BAK Forward: 5′-GTAGCCCAGGACACAGAGGA -3′, Reverse: 5′-ATAGCGTCGGTTGATGTCGT -3′; Noxa Forward: 5′-AGCTGGAAGTCGAGTGTGCT -3′, Reverse: 5′-TCCTGAGCAGAAGAGTTTGGA - 3′, Puma Forward: 5′-CCTGGAGGGTCCTGTACAATCT-3’, Reverse: 5′-TCTGTGGCCCCTGGGTAAG-3’ 3’, Actin Forward:5’-CACCAACTGGGACGACAT - 3’′, Reverse: 5′-ACAGCCTGGATAGCAACG-3’.

### Generation of RelA-YY1 Fusion construct

For generating the RelA-YY1 fusion construct, we first amplified human RelA from GFP-RelA plasmid (Addgene #23255) using primers harboring Kozak sequence at N-terminus and a mouse IgG2a hinge region at C terminus region. Hinge region sequence was according to pFuse-mouse-IgG2a. The resulted PCR product (RelA-Hinge) was cloned upstream to 3X-HA YY1 in pcDNA3.1 HA-YY1 plasmid (Addgene #104395) under CMV promoter resulting in the generation of RelA-hinge-3xHA YY1 fusion product. The RelA-hinge-3xHA YY1 fusion construct was further cloned into FUW lentiviral vector. Primers used were: Forward:5’-CATGGATCCGCCACCATGGACGAACTGTTCCCCCTCATC-3’ and Reverse: 5′-GTAGGATCCTGGGCATTTGCATGGAGGACAGGGCTTGATTGTGGGCCCTCTGG GGGA -3’.

### Cloning of Human Puma promoter in to Basic pGL2 Luciferase reporter vector

Primers were designed to clone human 3kb puma promoter in to Basic pGL2 Luciferase reporter vector. Human genomic DNA was used as a template, primestar GXL high fidelity enzyme and appropriately diluted primers were used for the amplification of 3kb Promoter. The amplified 3kb puma promoter was cloned into Kpn-1 and Nhe-1 sites of basic pGL2 luciferase reporter plasmid to generate pGL2-Puma reporter plasmid. Primers used to clone Human Puma Promoter were Forward:5’-GTAGGTACCCACCACTTACACACCATACACC -3’ and Reverse: 5′-GATTAATATAGCTAGCAGGCCGCCCGGCGGATCC-3’.

### Luciferase Reporter assay

P1234 3x-kB-L Luciferase reporter vector (Addgene#26699), FUW-RelA plasmid (subcloned from RelA-GFP plasmid), pcDNA3.1 HA-YY1 plasmid (Addgene #104395) were procured from addgene. pSV-β-Galactosidase Control Vector was purchased from Promega (Cat# E1081). Fusion construct plasmid was generated in the lab. Co-transfection of P1234 3x-KB-L luciferase reporter vector with indicated plasmids were performed with the help of transfection reagent Lipofectamine-2000 (Invitrogen Cat# 11668-019). 36 hours post transfection lysates were made in 1X Reporter lysis buffer (Promega Cat# E397A) containing protease inhibitor. Luciferase assay was performed by using luciferase assay substrate (Promega Cat# E151A) and β-galactosidase assay as an internal control for normalizing the transfection efficiency. 2-Nitrophenyl β-D-galactopyranoside (Sigma Cat#: N1127) was used as a substrate for β-galactosidase assay. 1M Sodium carbonate (SRL Cat#: 93857) was added to terminate the reaction. Similarly, Human 3KB puma promoter activity also analyzed in the presence of FUW-RelA and pcDNA3.1 HA-YY1 plasmid separately.

